# Embodied Intelligence Unlocks Autonomous Microscopy

**DOI:** 10.1101/2025.08.13.670210

**Authors:** Gang Huang, Zhengyang Zhang, Songlin Zhuang, Yang Wu, Zhihui Lu, Mingsi Tong, Huijun Gao

## Abstract

Advanced microscopy is a cornerstone of modern science, yet its potential is constrained by a reliance on manual operation, which struggles with the complexity and reproducibility of long-term experiments protocols. While scripting offers a degree of automation, it lacks the generalizability to adapt to new samples or dynamic biological events. A fundamental challenge is the absence of an intelligent system that can interpret high-level scientific intent and ground it in the physical actions of the microscope. To bridge this gap, we introduce an Embodied Intelligent Microscope System (EIMS) with hierarchical reasoning structure that reimagines the microscope autonomous control. This system leverages the advanced reasoning of large models to interpret complex user commands and decompose them into actionable steps. To solve the critical grounding problem, we constrain the model’s output to the feasible action space, effectively serving as the model’s “hands and eyes” in the physical world. We demonstrate that our system achieves zero-shot generalization on complex, multi-step protocols and successfully automates challenging biological missions requiring expert-level judgment, such as capturing the sparse spatiotemporal events of cell mitosis and locating scatteredly distributed organoids. This work establishes a new paradigm for scientific instrumentation—fusing high-level intent understanding with grounded, dynamic execution—and provides a generalizable framework for deploying embodied intelligence to accelerate autonomous scientific discovery.

## INTRODUCTION

Microscopy has revolutionized biomedical research, offering unprecedented windows into the intricate machinery of life. However, a significant semantic gap persists between a scientist’s high-level experimental goals and the low-level, tedious commands required to operate instruments^1,2^. This reliance on manual control not only limits the scale, speed, and reproducibility of experiments but also hinders the exploration of complex biological phenomena that unfold over long durations^3^. While script-based automation has addressed simple, repetitive workflows, these solutions are rigid and brittle; they lack the flexibility and adaptability to handle unforeseen events or the diverse requirements of discovery-driven science, necessitating bespoke programming for each new task^4,5^. For microscopy in particular, the sheer diversity of samples and scientific queries demands a level of flexibility and generalization that far exceeds the needs of other automated instruments^6,7^.

A critical bottleneck lies in grounding abstract scientific intent within the physical reality of the microscope^8^. Recent breakthroughs in artificial intelligence, particularly large models, offer transformative potential to bridge this gap with their remarkable capabilities in semantic understanding, reasoning, and planning. However, a significant weakness of these models is that they lack physical embodiment^9,10^, where they have no direct perceptual experience or intrinsic understanding of the consequences their actions have in the physical world. Furthermore, lacking knowledge of microscope operational procedures and relevant biological context, an ungrounded model might generate a plausible-sounding plan that is nonsensical or impossible for a specific physical system to execute. This, therefore, poses a pivotal challenge: to physically ground the abstract intelligence of large models within the microscope. The essential question is how these models can be embodied to autonomously translate scientific intent into diverse sample-specific experimental plans, and refine human intent when it conflicts with the operation constraints.

Here, we present an Embodied Intelligence Microscope System (EIMS) that reconceptualizes the microscope not as a passive imaging tool, but as an autonomous agent capable of closing the loop between intent, action, and perception. To solve the critical grounding problem, we introduce two core contributions. First, we designed a hierarchical agent framework where a high-level policy acts as a deliberative planner, interpreting complex intent and decomposing it into a sequence of subgoals. These subgoals are then executed by specialized, low-level executor modules that encapsulate the microscope’s physical capabilities and respond to real-time feedback. This structure grounds the model’s abstract plans in the microscope’s real-world affordances. Second, we developed a standardized model grounding protocol that makes the model aware of its embodiment by providing in-context information about available hardware, valid parameters, and the contextual logic for coordinating different tools, thereby constraining it to generate feasible and coherent task plans. This design has greatly enhanced the success rate of the model in complex tasks.

We systematically validated EIMS on a comprehensive benchmark spanning real-world biological applications, from tissue sections to live organoids, demonstrating performance that matches and even surpasses that of human experts. To prove its scalability, we successfully deployed the system on over 17 distinct microscopes peripheral devices, orchestrating complex, multi-device manipulations. Furthermore, through ablation studies, we compared different model architectures, including multimodal paradigms, to delineate a path forward for the field. Finally, we challenged EIMS with tasks demanding intelligent decision-making: the autonomous capture of spatially and temporally sparse biological events, such as recording sparse distributed organoids and tracking cell mitosis in real-time. The system’s success in these complex, dynamic tasks showcases its superior capabilities. Our work establishes a new paradigm for intelligent, autonomous microscopy. More broadly, it provides a generalizable framework for deploying embodied artificial intelligence in experimental science, poised to accelerate the pace of discovery.

## RESULTS

### An Embodied Architecture for Intelligence Microscopy

To address the challenges posed by intelligent microscopes (Fig. 1a), EIMS establishes a closed-loop framework integrating intention understanding, task decomposition, and dynamic execution through a hierarchical architecture (Fig. 1b and Fig. S1). Our objective is to learn a policy *π* that maps a complex observation *o_t_* — comprising the current visual field ( *I_t_* ), the microscope’s state vector ( *s_t_* ), and the natural language dialogue with the user ( *l_t_* )—to a sequence of executable actions, *A_t_* = [*a_t_*, *a_t_*_+1_,…, *a_t_*_+*n*_] . The policy is thus represented by the probability distribution *P*( *A_t_*∣*o_t_*) .

**Figure 1.**
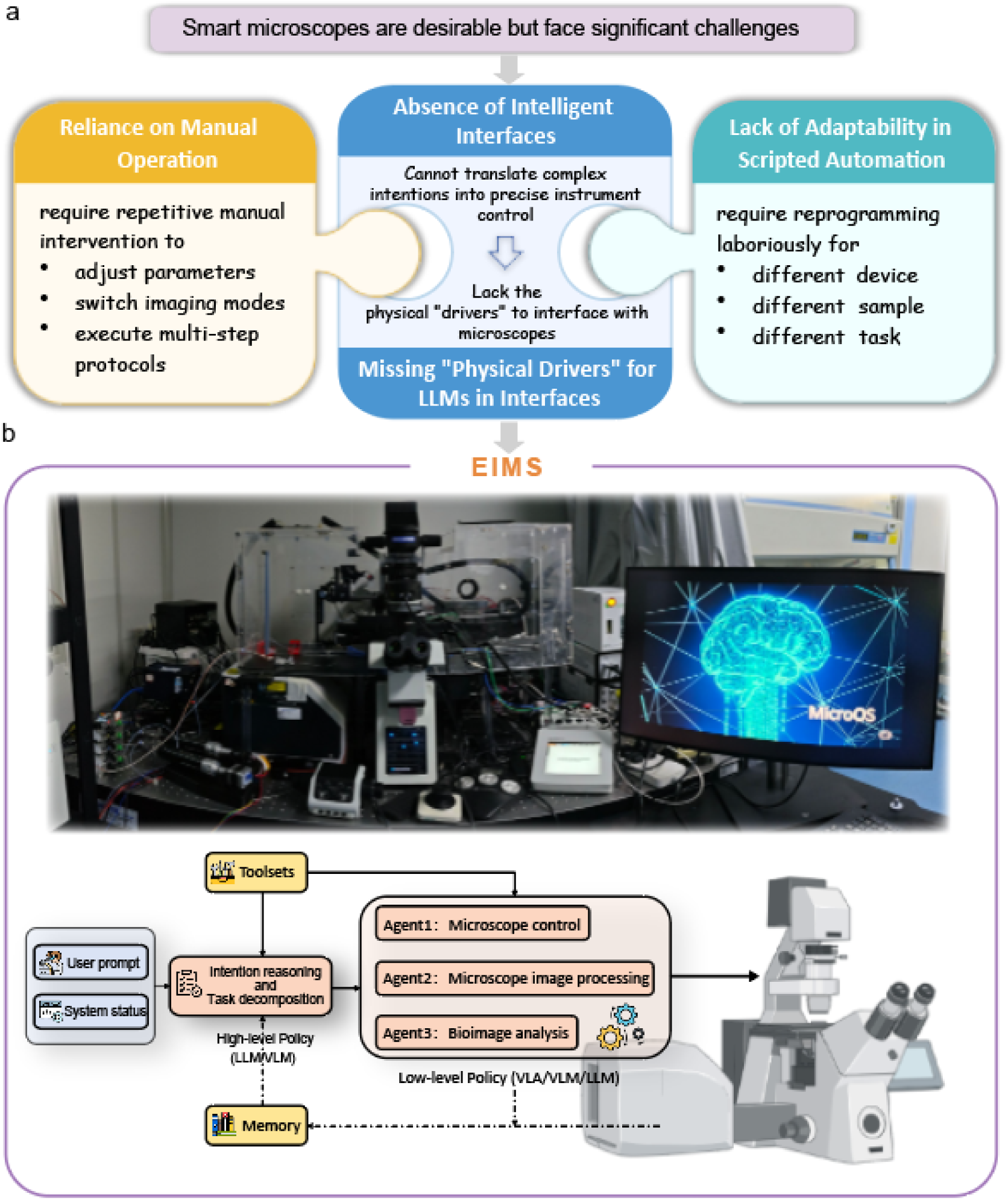
EIMS, an embodied intelligent system for autonomous microscopy. **a**, Fundamental challenges addressed by intelligent microscopy. **b,** The hardware platform and architectural overview of the EIMS framework that utilizes a hierarchical reasoning framework to connect intent with perception and action.

The fundamental challenge is bridging the gap between the abstract reasoning capabilities of large language models (LLMs) and the physical operation of complex scientific instruments like microscopes. The domain of microscopy is inherently characterized by immense procedural uncertainty. The manifold variations in biological samples—spanning different types, morphologies, and preparation methods—combined with the diverse imaging requirements of researchers, from rapid surveys to high-resolution three-dimension (3D) reconstructions, create a combinatorial explosion of potential action sequences for any given task. This makes predefining fixed workflows impractical.

Against this backdrop, the LLM’s inherent lack of physical grounding becomes a critical limitation. When presented with a high-level command such as “image this fluorescent tissue slide,” the model comprehends the semantic intent but responds with a generic, non-actionable narrative (e.g., “you can stain the samples, and then select the appropriate microscope for image acquisition and analysis.”). It cannot autonomously generate the precise, physically viable command sequence tailored to specific hardware.

To surmount this obstacle, we designed and implemented a hierarchical control policy. This architecture incorporates a high-level planning module responsible for decomposing the user’s ambiguous, high-level objective into a logically coherent sequence of intermediate subgoals. This decomposition process systematically reduces task complexity. Subsequently, a low-level execution module translates these concrete subgoals into atomic, executable commands for the microscope hardware. Our hierarchical architecture thus transforms an intractable decision-making problem into a structured, manageable workflow. It synergistically combines in-context learning to interpret user intent while imposing structural constraints that guarantee physically plausible outputs, thereby successfully solving the grounding problem among vast procedural uncertainty.

Specifically, the high-level policy, *_π_ ^h^* ,acts as the system’s cognitive core, responsible for long-horizon planning. It interprets abstract or ambiguous user instructions *l* and translates them into a structured, executable plan *l_π_* , a sequence of subgoals. This policy, *p^h^* (*l_π_*∣*l*) , must generate plans that adhere to the device’s capabilities, *L*_Π_ . Natural language is inherently ambiguous; a command like “find and snap a good cell” is subjective. To resolve this, our system initiates a clarifying dialogue and corrective interventions, modeled as a Bayesian filtering process to progressively refine its understanding of the user’s true intent. We treat the user’s ideal command as an unobservable latent state *x_t_* . The LLM’s generated plan at dialogue turn *t* serves as the observation *z_t_* , and the user’s feedback or additional information serves as the control input *u_t_* . Since LLMs can generate multiple possible instructions when given an ambiguous command, *z_t_* can be a probability distribution representing the LLM’s confidence in different instructions. By iteratively predicting and updating the posterior probability distribution of the true state, *p*(*x*∣*_t_ z*_1:*t* −1_ , *u*_1:*t* −1_ ) , we aim for the final model output to better estimate the user’s intended instruction.

Once a plan is finalized, the low-level policy, *π^l^* , takes over. It acts as the system’s motor controller, executing the plan. For each subgoal in *l_π_* , this policy, *p_l_* ( *A_t_*∣*o_t_* , *l_π_*) , generates concrete actions based on the current image *I_t_* and microscope state *s_t_* . In situations with multiple viable execution scheme, the system evaluates a set of possible affordances,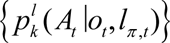. If the variance across these options is excessively high, indicating uncertainty, the system can trigger another perceptual cycle (e.g., acquire a new image, query the user) to reduce ambiguity until a high-confidence action can be selected.

While this hierarchical structure is not necessary for trivial tasks (e.g., “move Z-axis up 5 mm”), its power becomes evident in complex, multi-stage protocols that integrate image acquisition, processing, and analysis (Figs. 2 and 3). This architecture mirrors how human experts plan and execute large projects, dramatically improving success rates. Furthermore, it supports plug-and-play integration of new tools and ensures compatibility across diverse microscope platforms (Fig. 4), a critical advantage over rigid, platform-specific automation.

**Figure 2.**
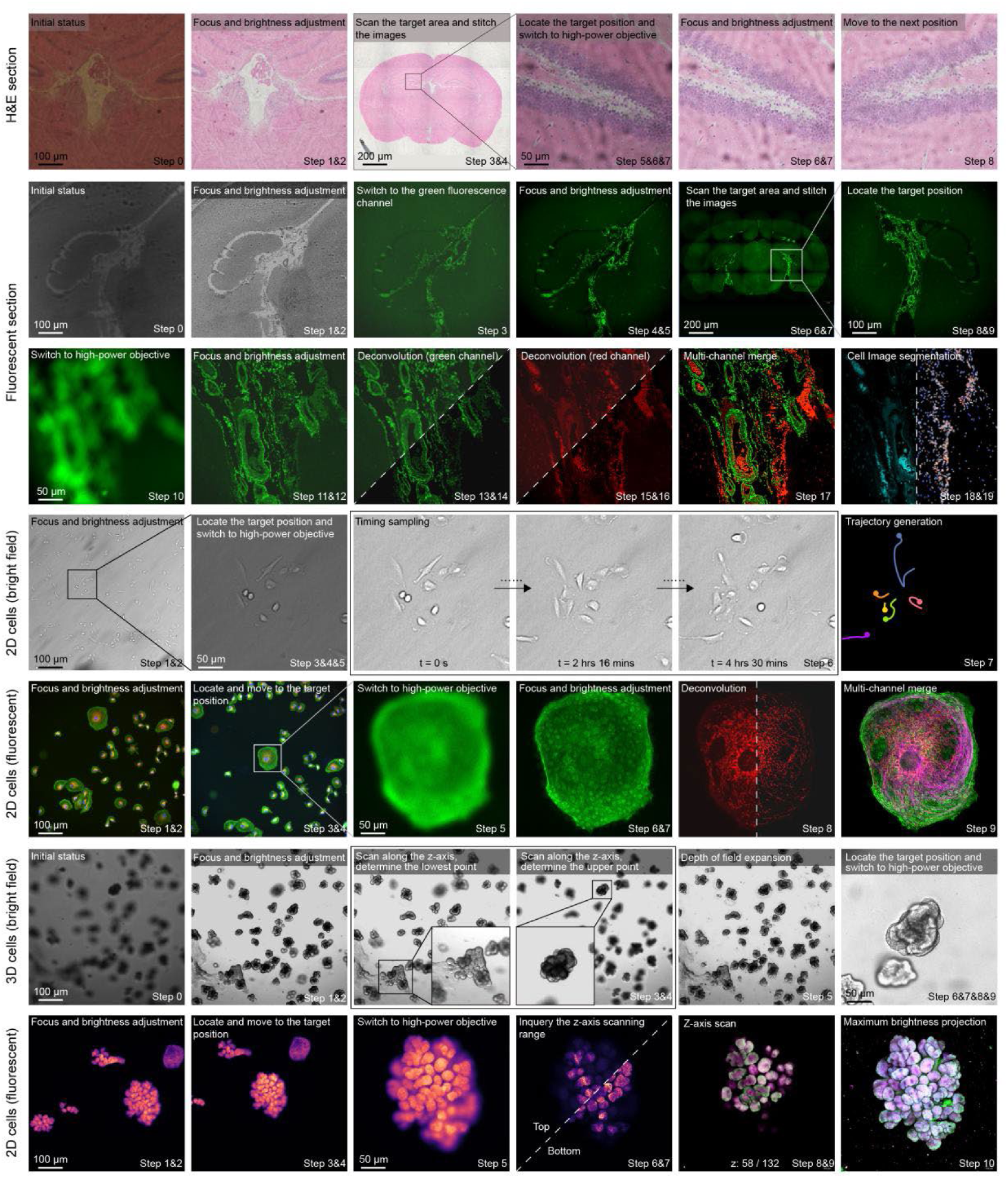
The core experimental workflow of the EIMS. Images of the operational steps of the EIMS, illustrating the automated experimental sequence and representative images acquired by the system, showcasing its adaptability to diverse sample types under various imaging conditions. The EIMS demonstrates robust performance across different sample preparations and imaging requirements, and integrates downstream tools for automated data post-processing and analysis.

**Figure 3.**
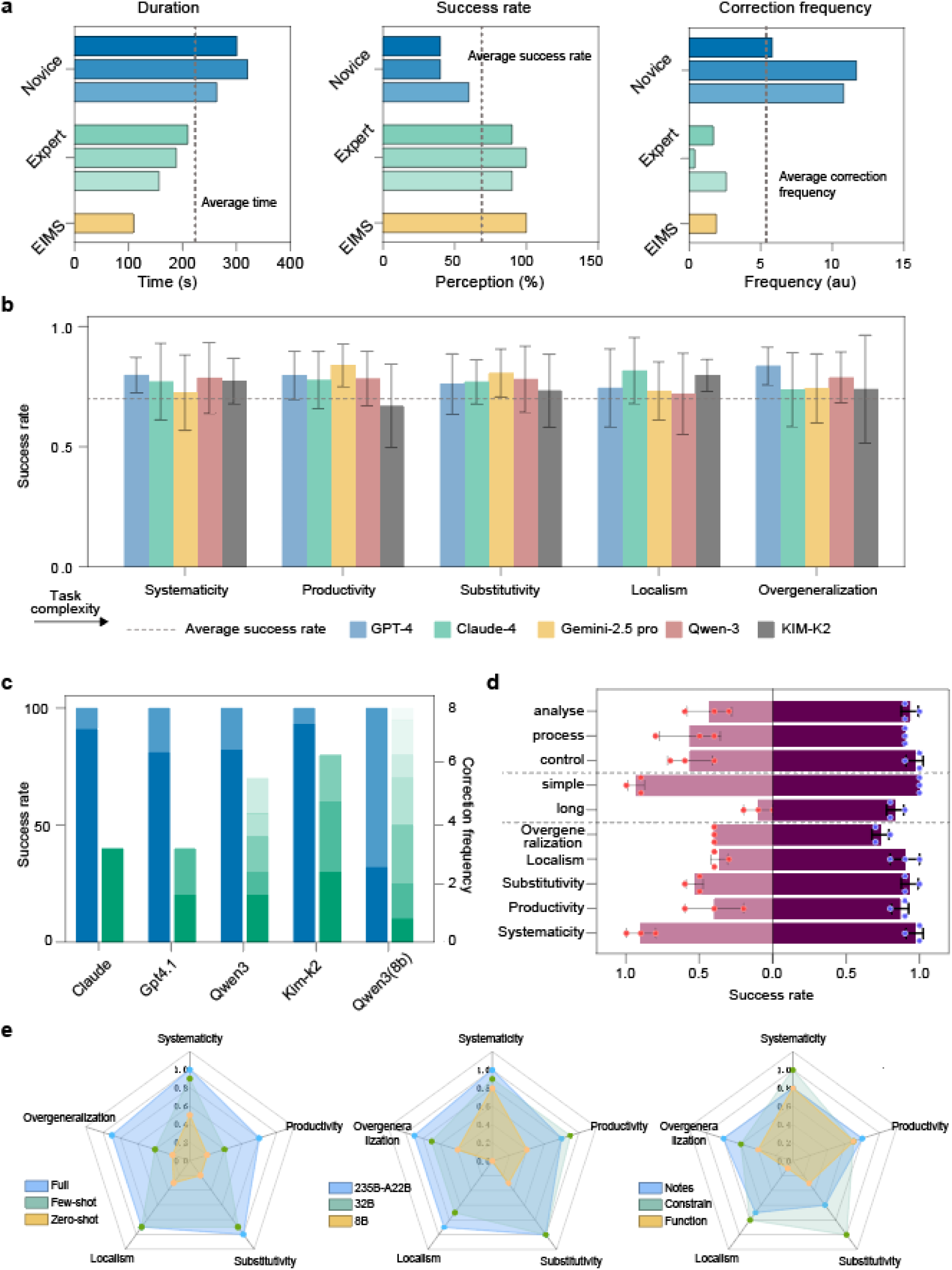
Performance evaluation of the EIMS. **a**, Quantitative comparison of MICROS with an expert human operator and a novice user. Performance was evaluated based on task completion time, success rate, and the frequency of corrective interventions required. **b,** Performance of MICROS across various generalization benchmarks when employing different foundation models. **c,** Ablation study showing task success rates and the corresponding number of corrections for different models with and without the correction module, demonstrating the module’s efficacy. **d,** Impact of the hierarchical architecture on success rates across different generalization evaluation sets. The analysis also includes performance on long-versus short-horizon tasks and on distinct sub-tasks: microscope control, image processing, and image analysis. **e,** Comprehensive analysis of model performance under various conditions. This includes a comparison of different tool-prompting strategies (function descriptions only, descriptions with single-tool examples, and descriptions with multi-tool chain-of-use examples); an evaluation of different model sizes against tasks of varying difficulty; and a comparison of model performance with different parameter settings on tasks of increasing complexity.

**Figure 4.**
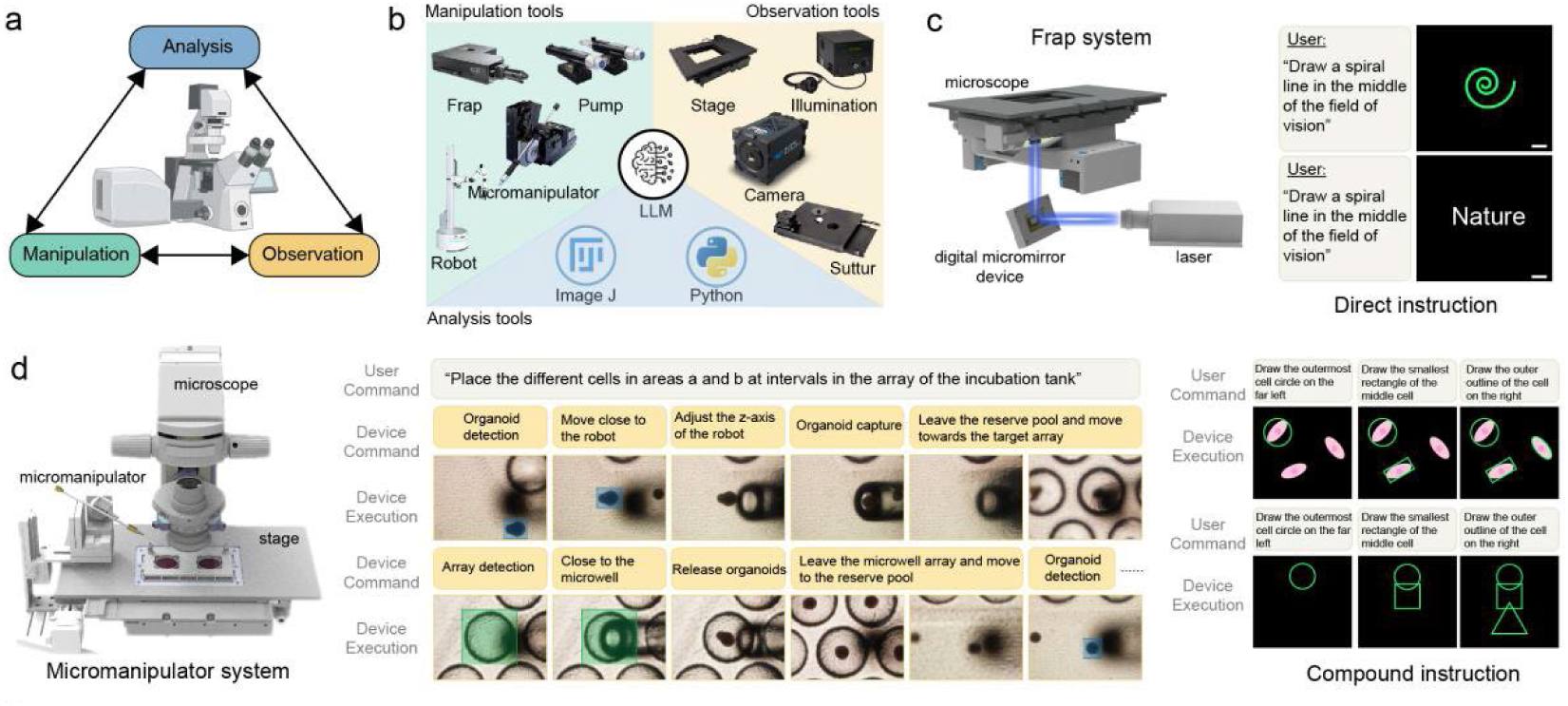
Scalability and cross-platform generalization of the EIMS framework. **a**, Schematic illustrating the paradigm shift in modern microscopy. Systems now integrate multiple tools to create a closed-loop workflow for the observation, manipulation, and analysis of biological samples. **b,** Schematic overview of the common hardware and software tools for observation, manipulation, and analysis in a typical automated microscopy system. **c,** Application of EIMS to control a microscope with an integrated FRAP module, demonstrating seamless cross-platform operation. **d,** EIMS controlling a micromanipulator to perform complex sorting of organoids under dynamic microscopic imaging, showcasing its capability for sophisticated automated biological manipulation.

### Performance evaluation of the EIMS

#### EIMS Performance Across Diverse Biological Applications

To evaluate EIMS performance, we established a comprehensive benchmark dataset encompassing a wide spectrum of biological samples and imaging conditions. This dataset, sourced from multiple biology experts and clinicians, includes tasks ranging from simple, single-step operations to complex, multi-stage protocols that require intricate physical manipulation and interactive user dialogue. Our analysis of the instruction dataset revealed two fundamental challenges to successful execution. First, natural language is inherently abstract, where a concise instruction of approximately 10 words can necessitate a sequence of 47 discrete action steps (Fig. S2). We quantified this information expansion, identifying an average token ratio of 1:72.5 between input commands and output actions. This substantial compression ratio introduces significant uncertainty into the model’s execution. Second, user instructions are typically goal-oriented (e.g., “assess the state of all organoids in the hydrogel droplet”) rather than procedural. This requires the model to not only interpret explicit commands but also to autonomously devise and execute the requisite methodologies and operational sequences to achieve the desired outcome.

The standard EIMS workflow commences with the high-level policy interpreting a user’s high-level command (Fig. S2a). Through an interactive loop, the system refines its understanding based on feedback, ultimately decomposing the high-level goal into a concrete, executable task plan. This plan is then passed to the low-level policy, where a suite of specialized agents orchestrates the execution. These agents directly control the microscope hardware, perform real-time image processing like denoising or super-resolution, and analyze image content to provide quantitative feedback to the user. All physical actions are constrained by predefined hardware limits, while the system leverages its embedded biological and optical knowledge to generate scientifically valid commands.

We systematically assessed EIMS on tasks categorized into 6 types based on sample nature and complexity, including bright-field and fluorescence imaging of tissue sections, two-dimension (2D) cell cultures, and 3D organoids. Across all categories, EIMS achieved a 100% success rate, defined as task completion with fewer than five corrective interventions (Table 1). Remarkably, 90% of all tasks were executed flawlessly without any need for correction. The small fraction of tasks requiring intervention (averaging just one correction per task) were consistently those with the highest complexity and a greater number of constituent steps. Representative examples of EIMS navigating these tasks are shown in Fig. 2, where it correctly selects appropriate objective lenses, switches between fluorescence channels, and adaptively adjusts exposure times and focus to optimize image quality. For complex multi-dimensional acquisitions, EIMS accurately interpreted user intent to configure the imaging parameters across space (x,y,z), time (t), and channels (c).

**Table 1.** EIMS Performance Metrics. This table presents the performance of EIMS across different sample model types, evaluated by success rates, number of corrections, and average step length. Each sample type comprises 10 distinct tasks.

|  | Success rate (%) | Average step | Correction frequency |
| --- | --- | --- | --- |
| Bright-field section | 100 | 8.5 | 1.7 |
| Fluorescent section | 100 | 10.7 | 2.4 |
| 2D cells (bright field) | 100 | 15.8 | 0.6 |
| 2D cells (fluorescent) | 100 | 19.8 | 1.8 |
| 3D cells (bright field) | 100 | 25.4 | 2.5 |
| 3D cells (fluorescent) | 100 | 26.8 | 3.4 |
| Average | 100 | 17.8 | 2.1 |

To contextualize these results, we benchmarked the performance of EIMS against that of a seasoned microscopy expert and a novice user. For this comparison, we selected long-horizon tasks from our benchmark suite. It is worth noting that the test experts did not contribute to the test set. As illustrated in Fig. 3a, both EIMS and the expert successfully completed all tasks with a minimal number of corrections. In contrast, the novice required frequent guidance, averaging over ten corrections or requests for assistance per task. This indicates that EIMS possesses a level of task understanding comparable to that of a human expert. Furthermore, EIMS demonstrated a significant efficiency advantage, completing tasks approximately 30% faster than the expert and 180% faster than the novice. These findings underscore the system’s consistency and robustness in interpreting user intent and recovering from procedural errors to successfully complete complex assignments.

#### Ablation Studies on Model Architecture and Design

The architecture of EIMS incorporates specific design choices to overcome the inherent challenges of grounding a language model in the physical world and lacking specialized microscopy manipulation knowledge. To dissect the contribution of each component, we performed a series of ablation studies using a purpose-built benchmark designed to assess generalization across five dimensions of increasing difficulty: Systematicity, Productivity, Substitutivity, Localism, and Overgeneralization (see Methods for more details). This tiered benchmark allows for a precisely dissect the model’s ability to generalize, from recombining known procedures to generating novel logic for unseen tasks.

We first evaluated the performance of different base models. To better reflect the differences among the various models, we temporarily hid the correction module. For the performance of different models, Four leading LLMs—Claude-4, GPT-4.1, Kimi-K2 and Qwen-3—were evaluated for their instruction interpretation, task decomposition and action execution accuracy. As shown in Fig. 3b, Claude-4 achieved a striking 96% success rate on the overall benchmark, outperforming GPT-4.1 (88%), Kimi-K2 (84%) and Qwen-3 (84%). All models performed near-perfectly on tasks requiring the recombination of existing primitives (Systematicity, Productivity, Substitutivity). However, their performance diverged on Localism and Overgeneralization tasks, which demand reasoning about novel situations. The error analysis revealed four primary failure situations: model hallucination (e.g., inventing nonexistent parameters), misinterpretation of user intent due to a lack of domain knowledge, procedural errors reflecting ignorance of standard microscopy workflows, and instruction forgetting in long-horizon tasks (Fig. S2). These findings highlight the challenge that the complexity of microscopy systems— stemming from the need to coordinate multiple device components and manage a vast range of samples and requirements—poses to current foundation models. Nevertheless, the introduction of an error-correction module elevated the success rate of all models to above 98%, with most tasks reaching completion after only two correction cycles.

Further ablation studies revealed the importance of our architectural and the setting of parameters in the model (Fig. 3d). We systematically omitted each design option to determine the contribution of each design. Replacing the hierarchical high-level/low-level policy structure with a single monolithic model caused a precipitous drop in performance, with the success rate on generalization tasks decreasing by 30%. We then assessed the components of our standardized control protocol. Providing examples of function usage (few-shot) dramatically improved performance over supplying only function definitions (zero-shot), underscoring the importance of demonstrating command-to-semantic mapping in microscopy control. However, this alone was insufficient for tasks requiring logical coherence, highlighting the critical need to also provide context on the coordinated use of different tools (+32.1% versus few-shot and +67.2% versus zero-shot). Furthermore, the inclusion of ‘Behavioral Constraints’ to standardize output formats and ‘Notes’ containing essential biological knowledge and operational principles further enhanced the success rate (+11.2% versus without Behavioral Constraints and +17.3% versus without Notes). We also observed a performance hierarchy linked to model size and architecture. Large Mixture-of-Experts (MoE) and dense models significantly outperformed smaller 8B-parameter models on complex reasoning. However, the introduction of the correction module substantially boosted the performance of these smaller models, increasing their success rate by 80%. Collectively, these results demonstrate that the cascaded architecture and the standardized control protocol enable EIMS to address abstract and complex tasks in microscopy control more effectively than non-specialized models.

#### Scalability and Cross-Platform Generalization

A foundational design principle of EIMS is its standardized hardware protocol, which confers cross-platform generalization and “plug-and-play” capability. Modern biological inquiry relies on an ecosystem of instruments where the microscope acts as a central hub for an observe-analyze-manipulate loop^11,12^, often integrated with a suite of ancillary devices (Fig. 4a-b). A truly intelligent operating system must orchestrate this entire hardware collective. To validate this capability, we successfully benchmarked EIMS on over 17 distinct devices commonly found in microscopy workflows, demonstrating its broad applicability and ease of adaptation to new hardware.

We conducted comprehensive benchmarking across diverse hardware setups and task complexities to evaluate the performance of EIMS. We assessed it on both simple and complex tasks derived from real-world user needs. For single-device tasks, such as executing a sequence of atomic commands, EIMS achieved a 100% command recognition and execution accuracy across all tested hardware. For complex benchmark, it includes multi-stage experiments and even involves cross-device coordination. For example, we use a micromanipulator to transfer heterogeneous organoids and arrange them into the desired pattern. This requires the simultaneous use of a micromanipulator, a motorized stage, and a camera, etc. EIMS demonstrated a profound ability to decompose high-level goals into synchronized sub-tasks, achieving a 96% full-pipeline success rate.

We highlight some examples that showcase the system’s emergent capabilities in complex tasks. First, we challenged the system to control a Fluorescence Recovery After Photobleaching (FRAP) system that can “drawing” precise patterns in the micro-scale with a laser. We tasked EIMS with interpreting abstract geometric descriptions in natural language. The system demonstrated remarkable spatial-geometric reasoning: it first used its perception module to identify target cells, then its planning module generated a series of 2D waypoints, which the laser followed to draw new shapes, precisely parsing dimensions and executing multi-step manipulations (Fig. 4c). Next, we escalated the complexity to a task where the microscope becomes an actor, not just an observer. We instructed EIMS to arrange 200 µm organoids into a specific pattern using a robotic micromanipulator (Fig. 4d). This required a symphony of coordinated control. EIMS orchestrated the motorized stage for coarse positioning and the micromanipulator for fine-grained placement. It used real-time visual feedback to detect the organoid, enabling a gentle, flexible grip, and translated the user’s linguistic description of the desired pattern into precise numerical coordinates for the micromanipulator. These experiments confirm that EIMS possesses the cross-platform generality to serve as the central nervous system for the automated laboratory of the future^13–15^.

#### Comparison with Alternative Embodied AI Architectures

We further evaluated multiple variants of EIMS, which mainly involved replacing the execution model with a multimodal model that can autonomously make decisions based on real-time microscopic images, as well as an end-to-end architecture with a direct-action execution policy. While EIMS paradigm uses an LLM to translate abstract goals into physical actions, itself is akin to an operator who is blindfolded and whose hands are tied, relying on external tools for all perception and action. This raises fundamental questions: What is the optimal cognitive architecture for an embodied microscope? Should intelligence be grounded primarily in language or in vision? And what advantages do end-to-end sensorimotor models offer? To explore this, we systematically compared three distinct model paradigms: the LLM-based approach of EIMS, a Vision-Language Model (VLM) that reasons based on real-time image input, and an end-to-end Vision-Language-Action (VLA) model that directly maps sensory input to motor actions (Fig. 5a-5c).

**Figure 5.**
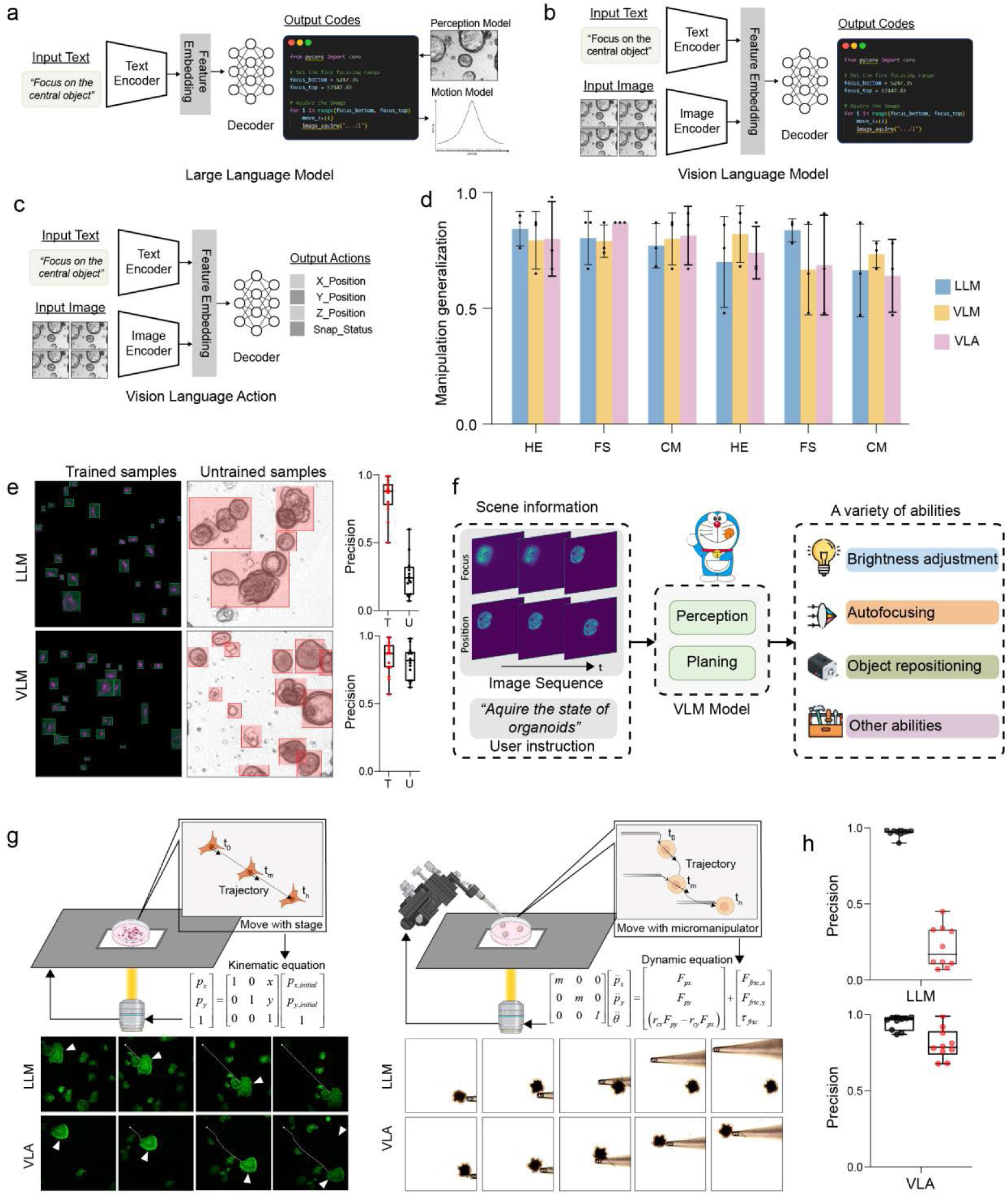
Comparison of EIMS performance with alternative architectural variants. **a–c**, Schematics of the evaluated EIMS variants, detailing the distinct input-output structures. The variants are powered by a Large Language Model (a), a Vision-Language Model (b), and a Vision-Language-Action model (c), respectively. **d,** Task performance of different model architectures in automated microscopy operations. **e,** Generalization comparison between the LLM and VLM variants across a range of different tasks. **f,** An illustration showing that the VLM’s ability to interpret visual data grants it superior task-generalization capabilities. **g,** Performance differences between the VLA and LLM variants, particularly highlighting the VLA’s effectiveness in tasks characterized by complex operational dynamics. **h,** In tasks with different complexities, VLA and LLM exhibit differences. The statistical graph represents the error in accuracy.

We designed a benchmark of real-time adjustment tasks (e.g., autofocusing, exposure adjustment) to quantify these differences. All models were tested in a zero-shot setting to probe their generalization capabilities. While all models could complete the tasks, their performance and failure modes were deeply illuminating (Fig. 5d). The LLM-based policy, equipped with specialized tools, achieved high precision. However, its success is fundamentally tethered to its visual perception and action-generation modules. A tool for assessing focus is useless for detecting objects; a detection model fine-tuned on one cell type may fail on another (Fig. 5e). This tool-based approach suffers from a lack of generalization. In contrast, VLMs, which internalize visual perception, demonstrated far greater flexibility. By grounding language in a native understanding of the visual world, a single VLM could be applied to multiple different tasks, effectively subsuming the function of many specialized tools (Fig. 5f).

The superiority of end-to-end integration became undeniable in tasks requiring complex dynamic control. For a basic action like moving a cell with stage from point A to B (Fig. 5g), a rule-based controller is sufficient, as the dynamics are linear: *x_t_*_+1_ = *x_t_* + *v*Δ*t* . But for a complex action like pushing a cell spheroid with the micromanipulator (Fig. 5h), the dynamics are highly non-linear, involving friction ( *F_friction_* ), fluidic drag ( *F_drag_* ), and contact forces ( *F_robot_* ):

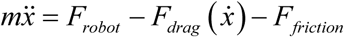

where *m* is the cell spheroid mass and *x* is the position. When the LLM-based agent attempted this using simple point-to-point commands, the final position error reached 72%. The VLA model, however, could continuously adjust its actions based on real-time visual feedback, mimicking human-like motor control and reducing the final error to just 7%. The results reveal a clear functional trade-off. VLA models excelled in tasks demanding highly adaptive and precise movements, which are indispensable for complex sample manipulation. However, for the primary role of a microscope — observation and image acquisition — the LLM-based architecture proved superior. Its strength lies in interpreting complex, language-based user intent and translating it into precise, multi-parameter instrument control, which is the most convenient and effective method for achieving high-quality scientific imaging results.

#### Performance of EIMS on samples with temporal and spatial sparsity

A defining feature of intelligence is not just executing plans, but the flexibility to adapt to new situations, react to unforeseen events, and respond to dynamic feedback^16^. Here, we demonstrate the advantage of EIMS over conventional automation: its ability to intelligently manage the stochasticity in long-horizen experiments. By fusing the reasoning of LLM with the principles of Event-Triggered Control (ETC), we transform the microscope from a passive recorder into an active decision-maker. In this paradigm, EIMS acts as an intelligent event detector and decision-making unit, autonomously determining the most scientifically valuable moments to act (Fig. 6a).

**Figure 6.**
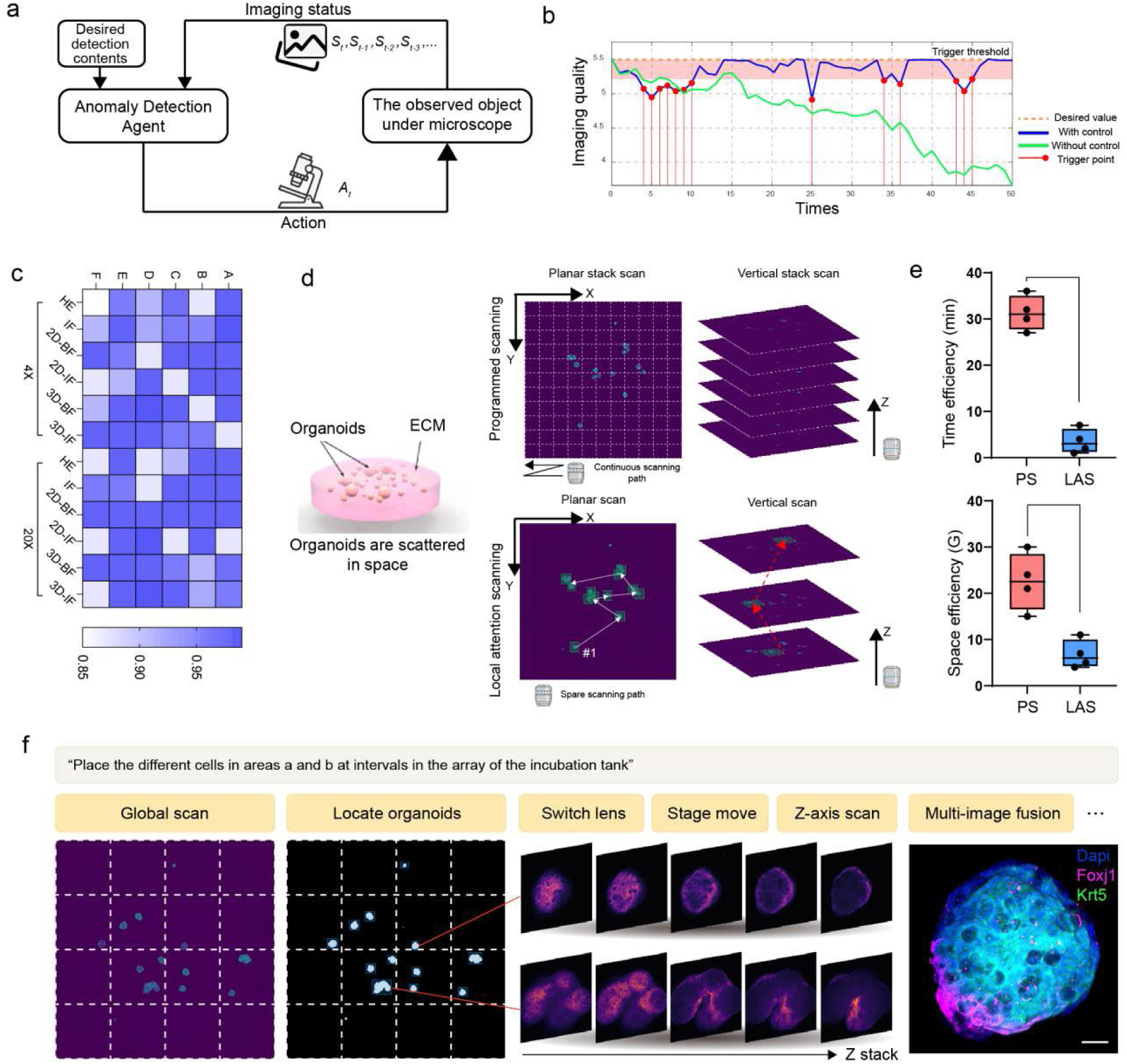
Performance of EIMS on samples with temporal and spatial sparsity. **a**, Schematic of the framework incorporating a state detection and correction module within the high-level policy. This policy not only responds to user commands but also autonomously analyses image quality to promptly correct imaging aberrations. **b,** Schematic demonstrating the integration of an event-triggered control strategy in EIMS. This enables robust, high-quality, long-term imaging of biological samples by effectively mitigating unknown perturbations. **c,** Success rates of the EIMS state detection and correction module across various sample types and imaging error categories. **d,** Schematic comparing conventional full-volume sampling with the EIMS local attention-based sampling strategy for spatially sparse samples. **e,** Comparison of acquisition time and data footprint between the different sampling strategies. **f,** Workflow of EIMS for imaging sparsely distributed organoids.

First, we tested the system’s ability to handle perturbations that would derail a conventional scripted experiment. Long-term live-cell imaging is a battle against entropy, where thermal drift, focus shifts, or sample contamination can slowly corrupt data quality. Manually programming for every “if-then” contingency is an intractable task. Instead, our goal is to develop an imaging program capable of sophisticated, human-like common-sense reasoning based on a comprehensive understanding of the real world, and then translate those inferences into actionable responses. To achieve this, we integrated EIMS with a Periodic Event-Triggered Control scheme (Fig. 6b).

For long-term imaging of biological samples, this can be conceptualized as a control problem under external disturbances, where our objective is to ensure that imaging quality converges to the desired value. The system’s quality is represented by a nonlinear dynamical model, governed by an intelligent control system that performs real-time image analysis to detect degradation events, such as sample drift or optical shifts. Upon detection of an anomaly, the system automatically triggers control commands to execute precise hardware adjustments, thereby restoring the system to its optimal state. To validate the efficacy of this closed-loop system, we introduced various perturbations to multiple biological specimens (Fig. S3). The system achieved a success rate of over 90% in event detection and compensation (Fig. 6c), ensuring robust long-term data acquisition. The minority of detection failures were primarily attributed to the precision limitations of the current visual encoder; specifically, the tokenization of image patches caused a loss of spatial resolution, rendering the model insensitive to specimen drift. We anticipate that targeted optimization of the model’s architecture will enhance its performance.

Beyond maintaining stability, this framework is powerful for capturing scientifically important but rare events. In this regard, we respectively demonstrate the model to tackle tasks involving temporal and spatial sparsity. For spatially sparse samples, like organoids randomly scattered in a hydrogel (Fig. 6d), if all the space is to be scanned equally, it will require an enormous number of resources (A full-volume scan of organoids in 24-well plates could take 29.54 hours and generate 1065.32 GB of data) (Fig. 6e). Instead, EIMS executes an event-triggered optimization strategy. It performs a rapid, low-magnification scan to identify spatial attention events— the locations of organoids—and then deploys targeted, high-resolution acquisition only on these regions (Fig. 6f). This intelligent decision-making process reduced imaging time by 2.57% (44.47 minutes) and data storage requirements by 2.39% (26.73 GB) while guaranteeing 100% coverage of the objects of interest.

The system’s intelligence is equally potent when confronting temporally sparse events. Biological processes are often punctuated by brief, dramatic transformations, such as cell mitosis, which are easily missed. The experimentalist faces a difficult trade-off: continuous high-speed imaging ensures capture but invites phototoxicity, while slow time-lapses conserve cell health but miss the action. EIMS resolves this dilemma by acting as an experienced biologist, by conducting sparse sampling to observe the current cell state, and then decide whether an interesting event has occurred and adjust the current acquisition parameters to achieve dynamic temporal resolution control for the capture of specific events. When imaging living cells, it maintained a patient, 5 minutes acquisition interval during the long, uneventful interphase. Upon detecting the subtle morphological cues preceding mitosis, the model triggered a temporal event, dynamically reconfiguring the system to a rapid, 30-second interval to capture the fleeting metaphase-to-anaphase transition with high fidelity (Fig. S4a). This adaptive strategy reduced cumulative phototoxicity by an average of 3.1-fold while increasing the capture rate of useful information by over 10-fold compared to non-adaptive methods (Fig. S4b).

We further combined spatial and temporal event-trigger capabilities to monitor thousands of cells simultaneously for mitosis events (Fig. S4c). The system uses a low-power scanning mode to patrol the entire cell population. Upon identifying multiple suspicious candidates based on their morphology, it dynamically allocates high-speed, high-magnification imaging resources exclusively to those cells. This allows for the simultaneous monitoring of ∼1300 cells at a slow interval, with imaging speeds in the selected regions of interest increasing by 200- to 5,000-fold faster as compared with the manual acquisition of timelapses in the whole investigated sample region (frame periods with the same acquisition settings: 134s (48-well plates), 512s (24-well plates), 2417s (12-well plates)). These experiments demonstrate the intelligent of EIMS, which can react and adapt in real-time. When the microscope itself knows what to image for, where to image, and when, the quality, throughput, and fundamental limits of scientific observation are dramatically redefined.

## DISCUSSION

In this work, we have demonstrated a fundamental paradigm shift in scientific instrumentation, moving beyond mere automation to true autonomy. The central challenge in advanced microscopy has been the immense gap between a scientist’s high-level, often abstract, intent and the low-level, tedious mechanics of instrument control. While robotics has recently undergone a profound transformation powered by large models, we asked if this revolution could be ported to the microscopic world. We answered this by introducing the principles of embodied artificial intelligence to microscopy for the first time, creating EIMS, a system that reimagines the microscope as a collaborative, intelligent agent. Our results show that this approach not only matches but in critical aspects surpasses human-level performance in complex, dynamic, and long-horizon experimental tasks.

The core challenge in achieving microscope autonomy lies in the practically infinite permutations of samples and imaging requirements, which generate an unenumerable space of operational procedures. This renders any approach based on fixed workflows brittle, as they inevitably fail when encountering scenarios outside their preprogrammed scripts. True intelligence is not the execution of a static script but the capacity to navigate an open world to find optimal solutions. Our investigation reveals that the key to this success lies in a hierarchical architecture that mirrors dual-process of human cognition^17^. The high-level policy engages in slow, methodical reasoning, interpreting complex user commands, formulating multi-step plans, and even engaging in clarifying dialogue to resolve ambiguity. This cognitive core then delegates subtasks to a reactive low-level policy that executes actions with speed and precision. This stratified design effectively solves the grounding problem; it gives the abstract reasoning of large models “hands and eyes” to interact with the physical world, constraining their vast knowledge to the practical real-world action. Our ablation studies validate the efficacy of this hierarchical design, which achieved a 100% task completion rate in our tests and demonstrated remarkable generalization and error-correction capabilities.

Furthermore, our work reveals that intelligent microscopy is not a matter of simple tool invocation. Unlike controlling a robotic arm for a singular task like grasping, operating a microscope requires orchestrating a complex system of interdependent components. Success demands an understanding not only of individual functions but also of the synergistic relationships governing their collective use. Indeed, our experiments prove that conventional tool-use paradigms, which merely provide function definitions, are largely ineffective in the microscopy context. By modeling the instrument as an integrated system of tools set, our approach boosted the task success rate threefold, confirming the critical importance of a holistic, systemic framework.

By comparing different AI paradigms, our work provides a clear roadmap for designing future intelligent instruments. We found that an LLM-based agent achieves high precision through its specialized modules but is consequently brittle, as its generalization is fundamentally limited by this predefined toolset. In contrast, a VLM natively integrates vision, enhancing perceptual generalization and allowing a single model to flexibly address diverse tasks without bespoke detectors. While current VLMs prioritize this flexibility, a promising future direction is the development of domain-specific VLMs optimized to rival the precision of tool-based methods. The VLA paradigm represents the ultimate integration—an operator who can see, act, and learn in a unified sensorimotor loop. Our micromanipulation experiments, where the VLA’s closed-loop control drastically reduced positioning error in a nonlinear dynamic task, conclusively show its superiority for complex physical interaction. This reveals a clear functional trade-off: VLA models are indispensable for tasks demanding adaptive, real-time physical manipulation. However, for the vast majority of microscopy applications centered on observation and image acquisition, the LLM-based architecture remains highly effective. Its strength lies in interpreting complex, language-based user intent and translating it into precise, multi-parameter instrument control, proving to be a robust and powerful paradigm for high-quality scientific imaging.

The implications of this work extend far beyond a single instrument, pointing toward a future of fully autonomous science^18–20^. Our successful integration of a wide range of instruments demonstrates a clear path toward closing the experimental loop, where an AI system could design a hypothesis, execute the experiment, analyze the data, and refine its next steps without human intervention. The task-aware imaging strategy we developed for sparse organoids is not a niche solution; it is a generalizable principle that could dramatically accelerate workflows in fields like digital pathology, where an AI could intelligently scan entire tissue slides at low resolution to find rare cancerous regions before performing targeted high-resolution analysis. Unlike previous data-driven microscopy methods trained for a handful of specific tasks, the near-infinite semantic understanding of large models unlocks a combinatorial explosion of capabilities, allowing scientists to command their instruments with the flexibility and creativity of natural language. The system’s ability to respond to unexpected perturbations and to intelligently capture rare spatiotemporal events, such as mitosis, proves it embodies the hallmark of true intelligence: flexibility.

Looking forward, the path is clear and exhilarating. We envision systems capable of continuous learning, where, much like a human collaborator, the AI’s proficiency and intuition grow with experience, requiring progressively less detailed instruction^21^. As the ecosystem of compatible hardware grows, developing sophisticated RAG systems or fine-tuning domain-specific models will become essential. The ultimate frontier lies in developing dedicated “world models” for the microscopic domain^22^. Such models would encode a deep, predictive understanding of micro-scale physics and cell biology, allowing the agent to simulate potential actions and choose the optimal plan, thereby achieving a profound level of adaptability. A tantalizing question this raises is how the real-world experiences gained by these embodied agents can be channeled back to improve the foundational models themselves, creating a virtuous cycle where acting in the world enhances an AI’s fundamental common-sense reasoning.

In conclusion, this work has laid the foundation for a new generation of scientific instruments. By endowing the microscope with the ability to understand intent, perceive its environment, and act purposefully within it, we have transformed it from a passive tool into an active partner in discovery. This fusion of embodied AI with experimental science does not seek to replace the researcher but to augment their capabilities, freeing them from the tyranny of manual operation to focus on the grand challenges of science. We have opened the door to a future where the speed of discovery is limited not by the dexterity of our hands, but by the breadth of our imagination.

## MATERIALS AND METHODS

### Data Collection and Test Set Construction

The training and evaluation dataset was compiled through extensive collaboration with multiple biology experts and clinicians. This dataset encompasses a wide spectrum of tasks, ranging from simple, single-step operations to complex, multi-stage protocols that demand intricate physical manipulation and interactive user dialogue. We stratified the data into six principal categories based on sample type: tissue sections, 2D cultured cells, and 3D cultured cells, with the brightfield and fluorescence imaging modalities. It is crucial to note that within each category exists a vast and diverse range of specific cell and tissue types, which are too numerous to exhaustively list. This intrinsic heterogeneity in sample morphology and experimental requirements presents a significant challenge to the model’s generalization capabilities.

To comprehensively evaluate the capabilities of the EIMS across its different modules, user requests within our dataset were structured to typically include three core components: (1) the desired image acquisition specifications, (2) the requisite image processing pipeline, and (3) the final analytical objectives. We quantified the complexity of the collected tasks by analyzing the number and distribution of procedural steps required for completion (Fig. S2). The mean number of steps per task was 23, with a minimum of 12 and a maximum of 37. Microscope control commands constituted the vast majority of these actions, accounting for approximately 82% of all steps.

### Generalization Dataset Design

To rigorously assess the model’s generalization, we systematically augmented and structured our collected data along five key dimensions: Systematicity, Productivity, Substitutivity, Localism, and Overgeneralization.

Systematicity evaluates the model’s ability to compositionally recombine learned functional primitives. For instance, after being trained on “acquire a Z-stack of the organoid using the 20x objective” and “scan the entire organoid hydrogel droplet with the 4x objective”, the model was tested with the novel command, “acquire a Z-stack of all organoids using the 4x objective”.

Productivity assesses the capacity to generate instruction sequences that are longer or more hierarchically complex than the training examples. This tests the model’s proficiency in handling long workflows and nested logic. Using the same training examples (“acquire a Z-stack of the organoid using the 20x objective” and “scan the entire organoid hydrogel droplet with the 4x objective”), a representative test command was “After scanning the entire organoid hydrogel droplet with the 4× objective, acquire a Z-stack of the centrally located organoid using the 20x objective”.

Substitutivity measures the model’s understanding of semantic equivalence by replacing terms with synonyms or functionally related concepts. This is critical for determining whether the model comprehends the underlying intent rather than merely memorizing command formats, and it probes its knowledge of biological and microscopy-specific terminology. For example, “image using the blue fluorescence channel” was substituted with “image the DAPI-labeled cells”.

Localism tests the ability to reuse a learned logical structure or concept in a new context, for a different objective, or with a different target. This evaluates whether the model understands the utility of its available tools. For example, having learned to “identify the organoid’s position with a low-magnification objective and then acquire a brightfield image with a high-magnification objective,” the model was tasked to “identify the organoid’s position with a low-magnification objective, then perform detailed data acquisition at that position using the blue and green fluorescence channels”.

Overgeneralization examines the model’s propensity to introduce novel API calls or programming constructs (e.g., autonomously generating a loop) not present in the training data. This characteristic is double-edged. On one hand, excessive unconstrained generation can lead to hallucinations and erroneous code that deviates from the user’s intent. On the other hand, this creative capacity enables the model to devise more efficient or novel solutions to complex tasks that were not explicitly demonstrated in the training data.

### Imaging Perturbations Dataset Design

To evaluate the model’s robustness, we constructed a dataset dedicated to common imaging perturbations. Data were collected across the six aforementioned sample categories under various challenging conditions. The primary perturbations tested were: axial (Z-axis) drift, motorized stage drift or abrupt sample movement, and variations in illumination intensity or photobleaching. We acquired data under both individual and combined perturbation scenarios. Furthermore, to assess generalization across different imaging scales, tests were conducted using multiple objective magnifications appropriate for each sample type. For instance, a 4× objective was deemed suitable as a low-magnification lens for organoids, whereas a 20× objective served a similar purpose for planar, adherent cells.

The evaluation of the model’s output correctness was based on a combination of expert annotations and quantitative metrics. For instance, we expected the model to reliably detect axial shifts exceeding two depths of field (DOF), identify illumination fluctuations greater than 10%, and recognize lateral drift corresponding to more than 10% of the field of view (FOV). Consequently, the absence of an alert when perturbations remained within these acceptable tolerance thresholds was also classified as a correct response.

### System Setup

The core of our experimental platform is an Olympus SpinSR spinning disk confocal microscope system, which features multiple programmable, motorized components essential for complex imaging protocols. The system includes: a tunable light source for brightness adjustment; a motorized stage for XY-plane sample positioning (Prior H117); a motorized Z-drive for focusing; a motorized sextuple revolving nosepiece equipped with 4×, 10×, 20×, 40×, 60×, and 100× objectives; an 8-position motorized filter wheel for selective observation of different fluorophores in widefield imaging; and a shutter to gate the widefield excitation path. For confocal imaging, the system is equipped with four solid-state lasers. Imaging is performed using a color camera (Olympus DP73), primarily for histopathological slides, and a high-sensitivity monochrome camera (Prime BSI) for fluorescence applications.

To validate the scalability and extensibility of our control algorithm, the system is further integrated with a suite of additional microscopy-related peripherals. This expanded setup includes a live-cell incubation chamber for time-lapse imaging, a microfluidics system for delivering cellular stimuli, a micromanipulation system for physical handling of biological samples, a FRAP unit for photomanipulation experiments, and a collaborative robotic arm for automated sample transfer. The micromanipulation subsystem itself comprise several motorized components, including the primary manipulator arm, a pneumatic pump for sample aspiration and holding, an oil-based microinjection pump, and a piezodrill for membrane penetration. Performance was also benchmarked on additional motorized stages, cameras, and laser systems to further demonstrate the platform’s modularity and extensibility.

### Scalability Test Dataset Design

For each piece of integrated hardware, we developed a dedicated test set based on common use-cases and operational protocols collected from experienced users. These test sets were structured into three tiers of increasing complexity: Basic Control Commands. Single-function instructions, such as “move the micromanipulator arm slightly to the left”; Composite Commands. Instructions requiring the coordinated action of multiple devices to achieve a systemic goal. For example, “Using the micromanipulator, sequentially transfer microspheres from the reservoir to the micro-array”. This command necessitates simultaneous control of the micromanipulator arm, motorized stage, and camera. Multi-Step Protocols. Tasks executed as a sequence of discrete commands, where the next command is issued only upon the successful completion of the previous one. The same microspheroid placing task, for instance, would be decomposed into: “Move to the reservoir area”, “Aspirate a microspheroid”, “Move to the array area” and “Release the microspheroid”, with each command being executed serially.

### Sample Preparation

#### Tissue Section Preparation

Mice brains werepost-fixed overnight in 4% PFA at 4°C, and subsequently cryoprotected by immersion in a 30% sucrose solution until equilibration. Tissues were then embedded in Optimal Cutting Temperature (OCT) compound (Tissue-Tek) and cryosectioned into 20 µm-thick coronal slices. For brightfield imaging (H&E Staining): Sections were air-dried and subsequently stained with Hematoxylin solution (Sigma-Aldrich) for 5 minutes, rinsed, differentiated in 0.3% acid alcohol, and then counterstained with Eosin Y solution (Sigma-Aldrich) for 2 minutes. Sections were dehydrated through a graded ethanol series, cleared with xylene, and mounted using a permanent mounting medium. For fluorescence imaging (Immunofluorescence): Sections were permeabilized with 0.3% Triton X-100 in PBS for 15 minutes and blocked with 5% normal goat serum (NGS) for 1 hour at room temperature. To visualize the vasculature, sections were incubated overnight at 4°C with a cocktail of primary antibodies: rabbit anti-alpha Smooth Muscle Actin (α-SMA, Abcam) to label arteries and rat anti-PECAM-1 (CD31, BD Biosciences) for endothelium. After washing, sections were incubated for 2 hours with species-specific secondary antibodies (Alexa Fluor 488 goat anti-rabbit and Alexa Fluor 594 goat anti-rat, Invitrogen). Finally, sections were counterstained with DAPI (4′,6-diamidino-2-phenylindole, 1 µg/mL) to label nuclei and mounted with ProLong Gold Antifade Mountant (Invitrogen).

#### 2D Cell Culture and Staining

A549 (human lung carcinoma) and COS-7 (monkey kidney fibroblast-like) cell lines were cultured in Dulbecco’s Modified Eagle’s Medium (DMEM, Gibco) supplemented with 10% fetal bovine serum (FBS) and 1% penicillin-streptomycin. Cells were maintained at 37°C in a humidified atmosphere of 5% CO₂. For imaging, cells were seeded onto glass-bottom µ-Dishes (ibidi) and grown to 70-80% confluency. For brightfield imaging: Live, adherent A549 cells were used directly. Prior to imaging, the culture medium was exchanged for phenol red-free imaging medium to reduce background. For fluorescence imaging: COS-7 cells were fixed with 4% PFA for 15 minutes, permeabilized with 0.1% Triton X-100 for 10 minutes, and blocked with 1% bovine serum albumin (BSA) in PBS. The actin cytoskeleton was stained with Alexa Fluor 488 Phalloidin (Invitrogen). Microtubules were labeled by incubating with a primary monoclonal antibody against α-tubulin (Sigma-Aldrich) for 1 hour, followed by a goat anti-mouse IgG secondary antibody conjugated to Alexa Fluor 594 (Invitrogen). Nuclei were counterstained with DAPI. Samples were maintained in PBS for imaging.

#### Mouse Airway Organoid Culture and Staining

Airway organoids were generated from primary mouse airway epithelial cells isolated from adult C57BL/6 mice, following established protocols. Briefly, isolated epithelial cells were suspended in Matrigel (Corning) and plated as droplets in 24-well plates. After polymerization, droplets were overlaid with organoid growth medium. Organoids were cultured for 10-14 days until maturation, with medium changes every 2 days. For brightfield imaging: live, mature organoids were imaged directly within their Matrigel matrix. For fluorescence imaging: mature organoids were released from Matrigel using Cell Recovery Solution (Corning), fixed in 4% PFA for 1 hour, and permeabilized with 0.5% Triton X-100. Whole organoids were blocked in 5% NGS and incubated overnight at 4°C with primary antibodies against Keratin 5 (KRT5, BioLegend) to mark basal cells and acetylated α-tubulin (Sigma-Aldrich) to mark ciliated cells. Following extensive washing, organoids were incubated with corresponding Alexa Fluor-conjugated secondary antibodies. Finally, organoids were counterstained with DAPI, mounted on glass slides using ProLong Gold Antifade Mountant, and imaged.

#### Statistical analysis

All the data are presented as the means ± SEMs unless otherwise stated. Microsoft Excel was used for statistical analysis. P < 0.05 was considered to indicate statistical significance.

